# Antibiotic-induced cell chaining triggers pneumococcal competence by reshaping quorum sensing to autocrine-like signaling

**DOI:** 10.1101/284125

**Authors:** Arnau Domenech, Jelle Slager, Jan-Willem Veening

## Abstract

*Streptococcus pneumoniae* can acquire antibiotic resistance by activation of competence and subsequent DNA uptake. Here, we demonstrate that aztreonam (ATM) and clavulanic acid (CLA) promote competence. We show that both compounds induce cell chain formation by targeting the D,D-carboxypeptidase PBP3. In support of the hypothesis that chain formation promotes competence, we demonstrate that an autolysin mutant (Δ*lytB*) is hypercompetent. Since competence is initiated by the binding of a small extracellular peptide (CSP) to a membrane-anchored receptor (ComD), we wondered if chain formation alters CSP diffusion kinetics. Indeed, the presence of ATM or CLA affects competence synchronization by shifting from global to local quorum sensing, as CSP is primarily retained to chained cells, rather than shared in a common pool. Importantly, autocrine-like signaling prolongs the time-window in which the population is able to transform. Together, these insights demonstrate the versatility of quorum sensing and highlight the importance of an accurate antibiotic prescription.

## Introduction

*Streptococcus pneumoniae* (the pneumococcus) is a Gram-positive diplococcus, member of the commensal microbiota of the human nasopharynx. However, the pneumococcus is also considered one of the leading bacterial causes of morbidity and mortality worldwide, being responsible for a wide variety of invasive and non-invasive diseases (O’Brien et al., 2009; Prina et al., 2015; Wahl et al., 2018). Pneumococcal infections are typically treated with antibiotics, which are considered a risk factor for the acquisition of resistant pneumococci in the nasopharynx. Actually, non-invasive infections (e.g. otitis media, non-bacteremic pneumonia or acute exacerbations of chronic respiratory diseases) are frequently caused by pneumococci with higher levels of antibiotic resistance than invasive strains (Kim et al., 2016; Kyaw et al., 2002). The colonization by antibiotic-resistant pneumococci in the human body can occur through several mechanisms, such as by the replacement of susceptible pneumococci by a resistant community-acquired strain, by spontaneous mutation prior to or during antibiotic therapy, or by the acquisition of a new antibiotic resistance allele from other pneumococci or closely-related Streptococci by the process of transformation (Schrag et al., 2000).

Transformation, defined as the uptake and assimilation of exogenous DNA, is an important mechanism of genome plasticity throughout evolutionary history, and is largely responsible for the rapid spread of antimicrobial resistance in the pneumococcus (Croucher et al., 2011). This process is regulated by competence (Figure 1A), a physiological state that involves about 10% of the pneumococcal genome (Aprianto et al., 2018; Claverys et al., 2009). Competence is induced by a classical two-component quorum sensing system in which the *comC*-encoded competence-stimulating peptide (CSP), is cleaved and exported by the membrane transporter ComAB to the extracellular space. CSP stimulates autophosphorylation of the membrane-bound histidine-kinase ComD, which subsequently activates the cognate response regulator ComE (Figure 1A) (Martin et al., 2013; Pestova et al., 1996). Upon a certain threshold CSP concentration, a positive feedback loop overcomes counteracting processes and the competent state is fully activated. One of the genes regulated by ComE, *comX*, encodes a sigma factor, which activates the genes required for DNA repair, DNA uptake, and transformation. CSP can be retained by producing cells (Prudhomme et al., 2016), but CSP also diffuses and can induce competence in neighboring cells (Christie et al., 2016; Håvarstein et al., 1995; Moreno-Gámez et al., 2017). Other environmental factors such as pH, oxygen, phosphate and diffusibility of the growth medium also influence competence development (Chen and Morrison, 1987; Claverys et al., 2002; Echenique et al., 2000; Yang et al., 2010). Thus, the initiation of competence can be considered as a combination of diffusion sensing and autocrine-like signaling (Doğaner et al., 2016; Moreno-Gámez et al., 2017).

The competent state is activated in response to several antibiotics, which thereby allow the bacterium to take up foreign DNA and potentially acquire antimicrobial resistance determinants (Prudhomme et al., 2006; Slager et al., 2014; Stevens et al., 2011). Hence, when antibiotic therapy is not appropriate or inadequate, competence activation of the commensal pneumococci resident in the nasopharynx can lead to the acquisition of resistant genes or virulence factors from its environment. Spread of antibiotic resistance is exacerbated by the fact that, coregulated with competence, *S. pneumoniae* expresses several bacterial killing factors, thereby using interbacterial predation to acquire foreign DNA (Kjos et al., 2016; Veening & Blokesch, 2017; Wholey et al., 2016).

We have shown previously that antimicrobials targeting DNA replication, such as fluoroquinolones, cause an increase in the copy number of genes proximal to the origin of replication (*oriC*) due to replication stalling (Slager et al., 2014). As the competence operons *comAB* and *comCDE* are located near *oriC*, these antibiotics induce competence. Aminoglycoside antibiotics such as kanamycin are thought to activate competence by causing the accumulation of misfolded proteins via mistranslation. Since these misfolded proteins are targeted by the HtrA chaperone/protease, the natural HtrA substrate CSP can accumulate and competence is activated (Stevens et al., 2011). While several classes of antibiotics have been tested for their ability to induce competence (Prudhomme et al., 2006; Slager et al., 2014), a systematic analysis of clinically relevant antibiotics and their effects on competence is lacking.

Here, we tested a panel of commonly prescribed antibiotics for their potential to induce competence. We found that the antibiotic aztreonam (ATM) and the beta-lactamase inhibitor clavulanic acid (CLA) induce competence. We show that both compounds bind to the non-essential D,D-carboxypeptidase PBP3. Consequently, cells are perturbed in their ability to separate, leading to the formation of long chains of cells. Cell chaining decreases diffusion of CSP into the extracellular milieu, thereby facilitating CSP’s interaction with membrane-bound ComD receptors on the producing cell itself and on daughter cells. This effectively changes the dynamics and shifts the major regulatory mode of competence from global quorum sensing to local quorum sensing, subsequently enhancing local competence induction and promoting horizontal gene transfer.

## Results

### Identification of clinically relevant antibiotics that induce competence

To monitor competence development, we utilized the ComX-dependent promoter P_*ssbB*_, driving expression of firefly luciferase (*luc*). We selected antibiotics on basis of their use for the treatment of several pneumococcal respiratory infections (otitis media, pneumonia or exacerbations of chronic respiratory diseases), as well as for the treatment of respiratory infections with other bacterial etiologies (Table S1). Cells of encapsulated strain D39V (Slager et al., 2018) were grown in C+Y medium at pH 7.3, a pH non-permissive for natural competence development under our experimental conditions (Moreno-Gámez et al., 2017), and antibiotics were added at concentrations below the minimum inhibitory concentration (MIC) to prevent large growth defects and cell killing. Only when antibiotics induce competence, the *ssbB* promoter is activated and firefly luciferase is produced. In line with previous reports, four antibiotics belonging to the fluoroquinolone and aminoglycoside classes of antibiotics robustly induced competence (Figure 1B)(Moreno-Gámez et al., 2017; Prudhomme et al., 2006; Slager et al., 2014; Stevens et al., 2011). Antibiotics from the macrolide and linezolid classes were not able to induce competence (Table S1). The beta-lactam subclass antibiotics, carbapenems and cephalosporins, also did not induce competence at any of the concentrations tested (Table S1). In contrast, the addition of aztreonam (ATM) and the combination of amoxicillin and clavulanic acid resulted in activation of P_*ssbB*-*luc*_. To test whether amoxicillin, clavulanic acid (CLA) or the combination of amoxicillin-CLA was responsible for competence induction, the compounds were also tested individually. Surprisingly, competence was not induced by the beta-lactam amoxicillin, but by clavulanic acid, an inhibitor of beta-lactamases. As the human nasopharynx is often colonized by non-typeable pneumococci, characterized by the absence of a polysaccharide capsule (Sá-Leão et al., 2006), we also tested whether ATM and CLA are able to induce competence in an unencapsulated derivative strain (strain ADP26). The deletion of the capsule did not affect competence induction by either of the drugs (Figure S1).

**Figure 1.**
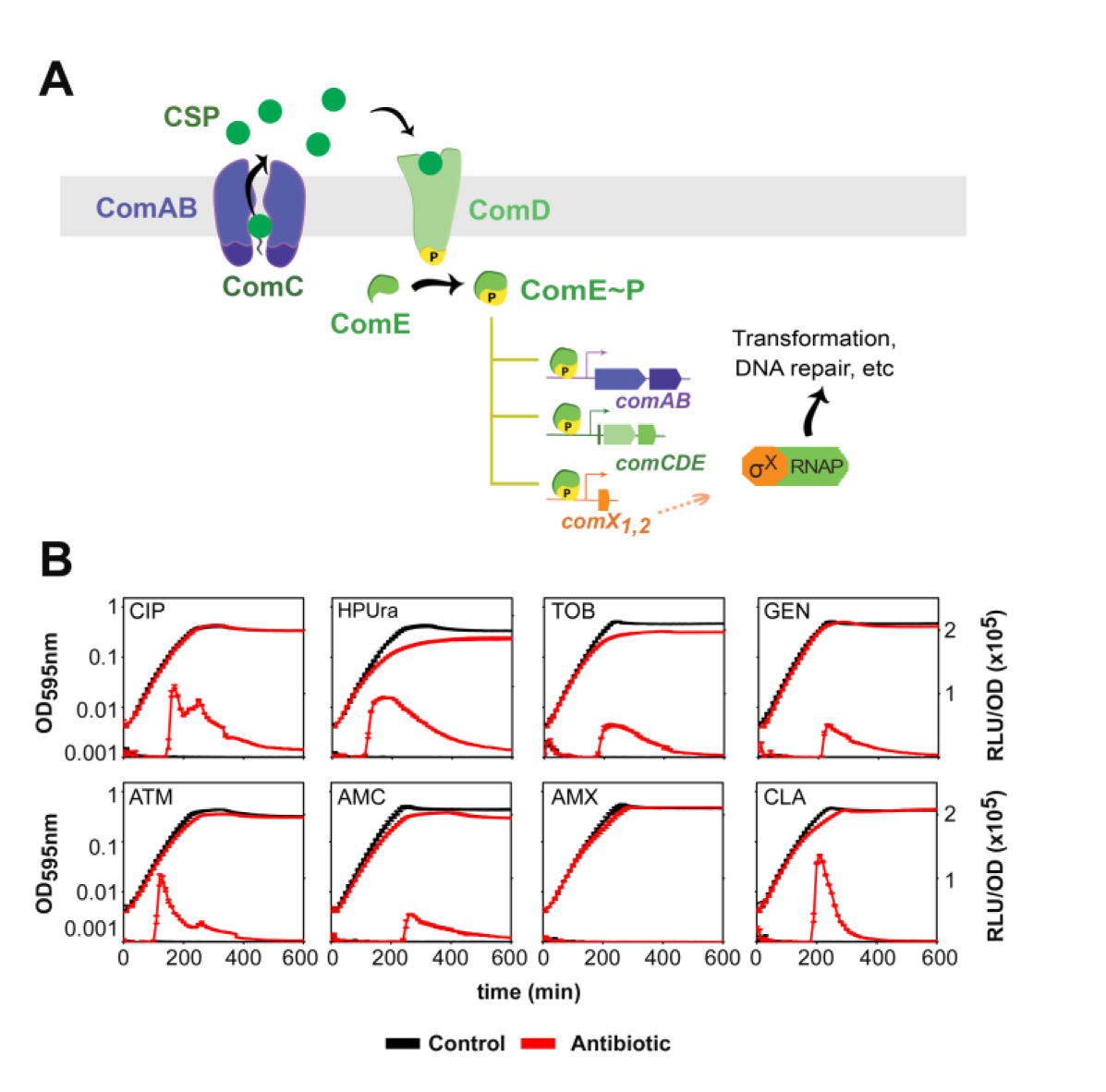
**A) Competence in *S. pneumoniae* is activated by several classes of antibiotics.** ComC binds the membrane protein complex ComAB and is processed and exported as CSP to the extracellular space. CSP then binds to the histidine kinase ComD, which is in the membrane as a dimer. Upon CSP binding, ComD autophosphorylates and transfers the phosphate group to the response regulator ComE (Martin et al., 2013; Pestova et al., 1996). The phosphorylated form of ComE (ComE~P) dimerizes and activates transcription of *comAB*, *comCDE* and *comX* by binding to their promoters (Håvarstein et al., 1995; Pestova et al., 1996). Synthesis of the alternative sigma factor ComX directs transcription of genes required for genetic transformation as well as other functions. **B) Growth curves (OD_595nm_) and bioluminescence activity (RLU/OD_595nm_) of *S. pneumoniae* in the presence of several antibiotics.** Strain DLA3 (*P_ssbB_*-*luc*) was grown in C+Y medium at pH 7.3, which is non-permissive for natural competence initiation, with (red lines) or without (black lines) addition of antibiotics. Average of 3 replicates and Standard Error of the Mean (SEM) are plotted. Concentrations of the antibiotics used: 0.4 μg/ml ciprofloxacin (CIP), 0.15 μg/ml HPUra, 28 μg/ml tobramycin (TOB), 10 μg/ml gentamicin (GEN), 28 μg/ml aztreonam (ATM), 0.12 μg/ml amoxicillin plus 2 μg/ml clavulanic acid (AMC), 0.12 μg/ml amoxicillin (AMX), and 2 μg/ml clavulanic acid (CLA). Induction of competence by ciprofloxacin and HPUra was shown before (Prudhomme et al., 2006; Slager et al., 2014).

Despite the introduction of conjugate vaccines, beta-lactam resistance is still a cause of concern in *S. pneumoniae.* To confirm whether ATM and CLA can induce competence in a strain with reduced susceptibility to beta-lactams, we tested a strain (ADP305) with a mutation in PBP2X (PBP2X^T550G^), which confers a MIC of 0.5 μg/ml and 0.64 μg/ml to penicillin G and cefotaxime, respectively. As shown in Figure S2, both antibiotics were still able to induce competence in this strain.

Together, this now extends the list of antibiotics capable of inducing competence to the following compounds: HPUra, mitomycin C, hydroxyurea, aminoglycosides, fluoroquinolones, trimethoprim, the beta-lactam aztreonam, and the inhibitor of beta-lactamases clavulanic acid.

### ATM and CLA promote horizontal gene transfer

To examine whether competence induction by ATM and CLA leads to increased horizontal gene transfer (HGT), we co-incubated two pneumococcal strains that are genetically identical except for a unique antibiotic resistance marker (tetracycline and kanamycin, respectively) integrated at different genomic locations. Since the extracellular pH is an important factor for natural competence development (Chen and Morrison, 1987; Moreno-Gámez et al., 2017; Prudhomme et al., 2006), we performed this experiment in two different growth conditions (pH 7.3 and pH 7.5, non-permissive and permissive conditions for natural competence, respectively) in presence or absence of ATM or CLA. As expected, at pH 7.3 no transformants were detected in the control condition. However, cells treated with either ATM or CLA showed significant HGT rates: (7.3 ± 2.3) · 10^-7^ and (2.3 ± 1.4) · 10^-7^, respectively. In addition, both ATM and CLA greatly enhanced HGT at pH 7.5 (Figure S3, Table S2).

ATM is mainly used to treat infections caused by Gram-negative bacteria as most Gram-positive bacteria, such as *S. pneumoniae*, are less susceptible to ATM. To test whether ATM could promote the transfer of DNA between a Gram-negative and *S. pneumoniae*, we co-incubated pneumococcal strain D39V with *Escherichia coli* strain DH5α. The *E. coli* strain used in this experiment carries a high-copy number plasmid, pLA18 (Slager et al., 2014), containing a tetracycline-resistance allele flanked by homology regions with the non-essential pneumococcal *bgaA* locus. At 28 μg/ml of ATM, *E. coli* is readily lysed while competence is induced in *S. pneumoniae* (Figure 1B). Importantly, a high fraction of *S. pneumoniae* transformants with the integration plasmid was observed, demonstrating that ATM not only promotes competence, but can also enhance DNA transfer by killing ATM-susceptible donors (Table S3).

### ATM and CLA do not induce competence via HtrA or altering gene dosage

So far, two different molecular mechanisms of competence induction by antibiotics have been described. The first mechanism is via substrate competition of the HtrA protease, which degrades both CSP and misfolded proteins (Cassone et al., 2012; Stevens et al., 2011). An increase of misfolded proteins induced by aminoglycosides would induce competence, because HtrA is occupied and CSP degradation is reduced. The second mechanism described applies to antibiotics that stall DNA replication elongation such as fluoroquinolones or HPUra. These drugs cause replication forks to stall, while DNA replication initiation continues, resulting in an increase in copy numbers of genes close to the origin of replication (including both early competence operons *comAB* and *comCDE*) (Slager et al., 2014).

We confirmed that strain ADP309, carrying a mutation in *htrA* that renders the catalytic domain inactive (HtrA^S234A^), is hypercompetent compared with the wild type (Figure S4; Stevens et al., 2011). However, competence was still induced in this strain by ATM and CLA, as well as by the aminoglycosides gentamycin and tobramycin.

To test whether ATM and CLA induce competence via altering the gene dosage of the early competence operons, we performed marker frequency analysis. As shown in Figure 2A, a shift in origin to terminus ratio was observed after the addition of HPUra; however, the presence of ATM or CLA did not lead to an increase of the *oriC*-*ter* ratio. To uncover potential transcriptional changes upon ATM or CLA treatment, we performed transcriptome profiling using DNA microarrays. We analyzed the rapid (15 minutes after addition) and adaptive (cells growing with the compound) transcriptional responses to ATM and CLA. Experiments were performed using a *comC* mutant strain to prevent the activation of competence, which will obscure data analysis. These analyses validated the marker frequency experiments and no differential gene expression of origin-proximal genes was observed (Figure 2B). Furthermore, both compounds, at competence-inducing concentrations, had minor effects on the global transcriptome, suggesting that their effects are on the post-transcriptional level (Tables S4 and S5).

**Figure 2.**
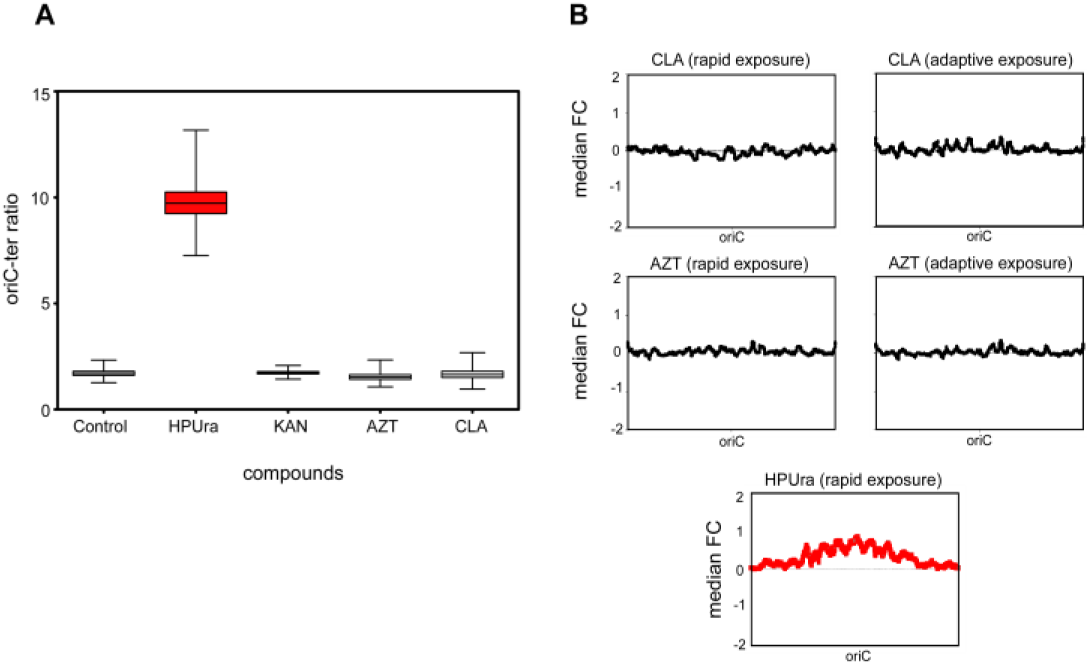
**A) Effect of antibiotic treatment on origin-terminus ratio.** Boxplots represent *oriC*-*ter* ratios as determined by real-time qPCR. Whiskers represent the 10^th^ and 90^th^ percentile of data from Monte Carlo simulations. Strain DLA3 (*P_ssbB_*-*luc*) was grown in medium without (control) or with the following compounds: 0.15 μg/ml HPUra, 28 μg/ml kanamycin (KAN), 28 μg/ml of aztreonam (ATM) and 2 μg/ml of clavulanic acid (CLA). Red box (HPura) matches with previous data showing an increase of the *oriC-ter* ratio (Slager et al., 2014). **B) Transcript copy number changes.** Every point is the median fold-change of 51 genes as a function of the central gene’s position. Both ATM and CLA do not affect the *oriC*-*ter* ratio. HPUra analysis from Slager et al. is shown in red, as a positive control of *oriC*-*ter* ratio shift (Slager et al., 2014).

### ATM and CLA target PBP3 and induce cell chaining

It is well known that both ATM and CLA have an impact on cell wall synthesis. Specifically, it has been shown that they can directly interact with PBP3 (Kocaoglu et al., 2015; Severin et al., 1997). To assess whether perturbing cell wall synthesis could lead to activation of competence, we employed CRISPR interference (CRISPRi), allowing us to downregulate essential genes involved in cell wall biosynthesis (Liu et al., 2017). Competence development was not influenced by downregulation of either genes involved in peptidoglycan precursor synthesis (*murA*-*F*, Figures S5-S6) or genes encoding class B PBPs (transpeptidase only) *pbp2b* and *pbp2x.* However, when the genes encoding class A (dual transglycosylase and transpeptidase) PBP1A, or the D,D-carboxypeptidase PBP3 were repressed using CRISPRi, competence was strongly induced under otherwise non-permissive conditions (Figure 3A).

**Figure 3.**
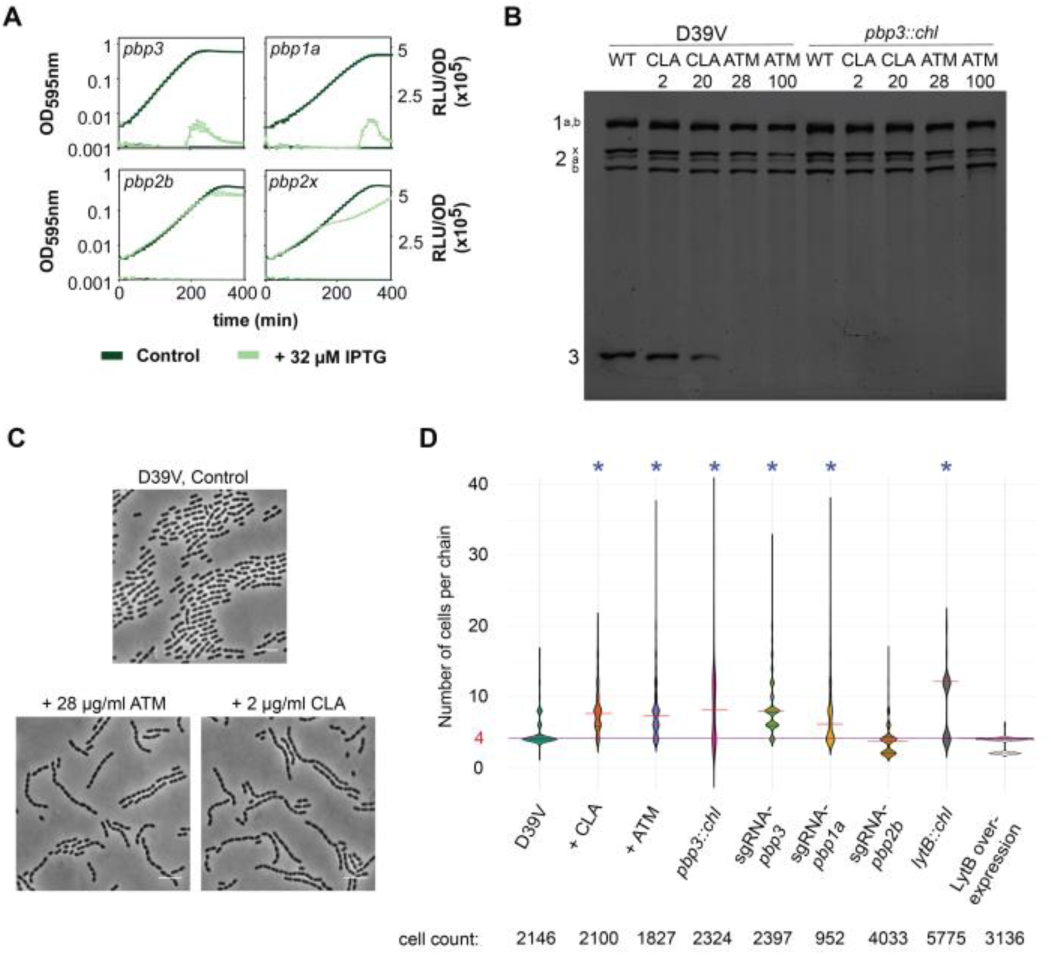
Aztreonam and clavulanic acid induce cell chaining by binding to PBP3. **A) CRISPRi-dependent downregulation of *pbp1a* and *pbp3* leads to competence induction.** Depletion of *pbp1a* and *pbp3* by induction of dCas9 with IPTG upregulates competence. In contrast, depletion of *pbp2b* and *pbp2x* does not have any effect on the regulation of this process. Detection of competence development was performed in C+Y medium at a non-permissive pH (pH 7.3). 32 μM IPTG was added to the medium at the beginning. The values represent averages of three replicates with SEM. **B) Representation of the PBP profiles of whole cells.** D39V and a Δ*pbp3* were treated with 2 μg/ml and 20 μg/ml of clavulanic acid (CLA), and 28 μg/ml and 100 μg/ml of aztreonam (ATM) and subsequently labeled with Bocillin-FL. Both ATM and CLA bind PBP3 in the D39V strain. Numbers indicate the different PBPs (i.e. 2b = PBP2B, 3 = PBP3). **C) ATM and CLA induce chain formation.** Phase-contrast images. Cells were grown in C+Y pH 6.8 until OD_595nm_ 0.4 (stationary phase). Scale: 6 μm. **D) Length of the chains.** The horizontal red line indicates the average of the number of cells per chain, while the purple line represents the control condition. The addition of ATM or CLA results in the formation of longer chains, as does the deletion and depletion of *pbp3*and the deletion of *pbp1a.* In contrast, *pbp2b* depletion does not lead to a chain-forming phenotype. The absence of *lytB* also resulted in an increase of chain length, while its complementation restores normal chain length.

To corroborate that *pbp2b* and *pbp2x* do not upregulate competence, we repeated the same experiment in a permissive pH for natural competence. As expected, *pbp1a* and *pbp3* repression resulted in a stronger induction of competence, while repression of *pbp2b* and *pbp2x* did not influence competence development (Figure S7).

To confirm that ATM and CLA bind PBP1A and/or PBP3, we used fluorescently labelled Bocillin (Bocillin-FL). As shown before (Kocaoglu et al., 2015; Severin et al., 1997), ATM and CLA bind PBP3, with ATM having a higher affinity to PBP3 than CLA (Figure 3B). As we were not able to clearly separate PBP1A and PBP1B, we cannot conclude whether ATM and/or CLA also bind to one of these PBPs. Since *pbp3* is not essential (Liu et al., 2017), we constructed a deletion mutant. In line with the CRISPRi results, the Δ*pbp3* (strain ADP30) displayed hypercompetent behavior (Figure S8). Importantly, ATM and CLA do not further induce competence in the Δ*pbp3*, indicating that PBP3 is the main target of these compounds (Figure S8).

To examine the effects of ATM and CLA and the downregulation of *pbp1a* and *pbp3* on cell morphology, we performed microscopy analysis on exponentially growing cells (OD_595nm_ 0.1), when the wild-type D39V strain becomes naturally competent at pH 7.9 (Moreno-Gámez et al., 2017). In contrast to cells with downregulated *pbp2b* or *pbp2x* (Berg et al., 2013; Land et al., 2013; Liu et al., 2017; Peters et al., 2014), individual cell size and morphology were only slightly affected by ATM, CLA or *pbp1a* and *pbp3* perturbation. However, in all cases, pneumococci formed longer chains of unseparated cells (Figure S9). When cells were grown until stationary phase (OD_595nm_ 0.4), chain formation was even more evident (Figure 3D).

Other beta-lactams, such as amoxicillin, ampicillin, piperacillin or cefotaxime, also have a strong affinity for PBP3, but also for PBP2X (Kocaoglu et al., 2015). The inactivation of PBP2X seems to counteract the effect of PBP3 depletion, because these drugs did not induce chain formation (Figure S10). These results suggest that cell chaining by ATM and CLA could be responsible for competence induction. To test this hypothesis, we performed a multi-dose checkerboard experiment with 8 different concentrations of both ATM and CLA (Figure S11). Indeed, the effects of ATM and CLA on competence activation are additive, until a certain maximum effect size, likely corresponding to the maximum chaining capacity.

### Cell chaining is responsible for ATM- and CLA-induced competence

To test whether ATM and CLA induce competence by specific binding to PBP3 or because of cell chaining, we generated a knockout of the gene encoding the major autolysin LytB (strain ADP21). LytB mutants are well known to form chains due to their lack in muralytic activity at cell poles (De las Rivas et al., 2002; García et al., 1999; Rico-Lastres et al., 2015). In line with the hypothesis that cell chaining induces competence, the Δ*lytB* mutant showed a hypercompetent phenotype, and readily developed competence even at pH 7.3, at which wild type cells do not become naturally competent (Figure S12A). Importantly, complementation by ectopic expression of LytB in the Δ*lytB* (ADP43), restored the normal diplococcus phenotype and restored competence development to wild-type-like (Figures S9 and S12B).

Finally, to test whether ATM or CLA induction is lost in the Δ*lytB* mutant, we have tested the effect of ATM and CLA in the Δ*lytB* and the complementation strain (Figure S13). In the absence of IPTG (chaining phenotype, Figure S13A), this strain is naturally hypercompetent, compared with the wild type DLA3, which did not become naturally competent at this pH. Under these conditions, ATM and CLA can only slightly accelerate competence development, relative to the control condition. LytB complementation by the addition of IPTG in ADP43 restores the normal phenotype and, as a result, the strain behaves as DLA3, confirming the role of chain formation in the regulation of competence (Figure S13B).

Besides Δ*pbp3* and Δ*lytB* mutants, we also tested competence induction in a Δ*divIVA* mutant, which also displays extensive chaining. In line with our hypothesis, this mutant has a hypercompetent phenotype as well (Figure S14). However, it should be noted that the deletion of *divIVA* has a strongly pleiotropic effect (Fadda et al., 2007), and competence development might be affected through more than one mechanism in this strain.

### Cell chaining modifies quorum sensing into autocrine-like signaling

We hypothesized that antibiotic-induced chaining reduces diffusion of CSP, thus altering synchronization of competence within the population. Under our standard plate-reader conditions (at the population level), encapsulated *S. pneumoniae* D39V cells release CSP into the medium, and when the CSP concentration reaches a critical threshold, cells activate competence in a synchronized manner (Moreno-Gámez et al., 2017). This results in a very steep RLU (Relative Luminescence Units) slope from the P_*ssbB*_-*luc* reporter at different inoculation densities (Figure 4, green line). In contrast, in the presence of ATM or CLA, the RLU increase for lower inoculation densities starts earlier, since both compounds induce competence; however, the slope of light production is less steep, indicating reduced synchronization of competence throughout the population (Figure 4).

**Figure 4.**
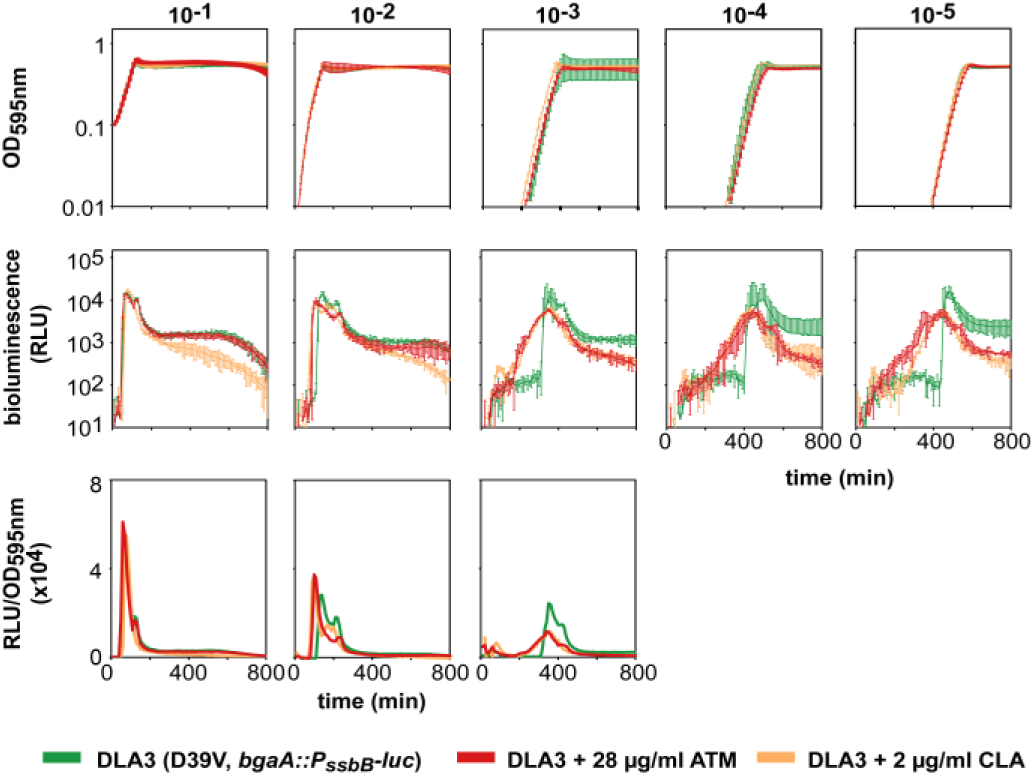
Synchronization of competence is affected by chain formation. Cells were grown in C+Y at competence-permissive pH 7.6, without antibiotics (green lines), in the presence of 2 μg/ml clavulanic acid (orange lines) or with 28 μg/ml of aztreonam (red lines). At pH 7.6, cells become naturally competent but with a certain delay compared to pH 7.9. Therefore, the inducing effect of ATM and CLA can be more easily visualized at this pH. Average of 3 replicates and Standard Error of the Mean (SEM) are plotted for each of five inoculation densities: OD_595nm_ 10^-1^, 10^-2^, 10^-3^, 10^-4^ and 10^-5^. RLU/OD could not be accurately calculated for the two lowest inoculation densities, due to the OD detection limit of our plate reader.

### Single-cell analysis of competence activation and signal propagation

For a better understanding of how competence is initiated and spread at the single-cell level in the wild-type population, we employed fluorescence microscopy and flow cytometry experiments. To first establish whether CSP produced by our wild-type D39V strain is shared in a common pool, explaining the rapid synchronization of the population, we repeated a previously published time-lapse microscopy experiment and grew wild-type pneumococci expressing a SsbB-GFP fusion (Aprianto et al., 2016) together with a Δ*comC* mutant strain that also contains the SsbB-GFP fusion and constitutively expresses a cytoplasmic RFP (Moreno-Gámez et al., 2017). We observed that wild-type cells became competent after 80 minutes as shown by the expression of SsbB-GFP, and the Δ*comC* cells start to express SsbB-GFP at the same time-frame, independent of whether cells touch each other or not (Figure 5A). This validates our assumption that wild-type cells share CSP in a common pool and can trigger competence in neighboring cells, without the necessity of direct cell contact.

**Figure 5.**
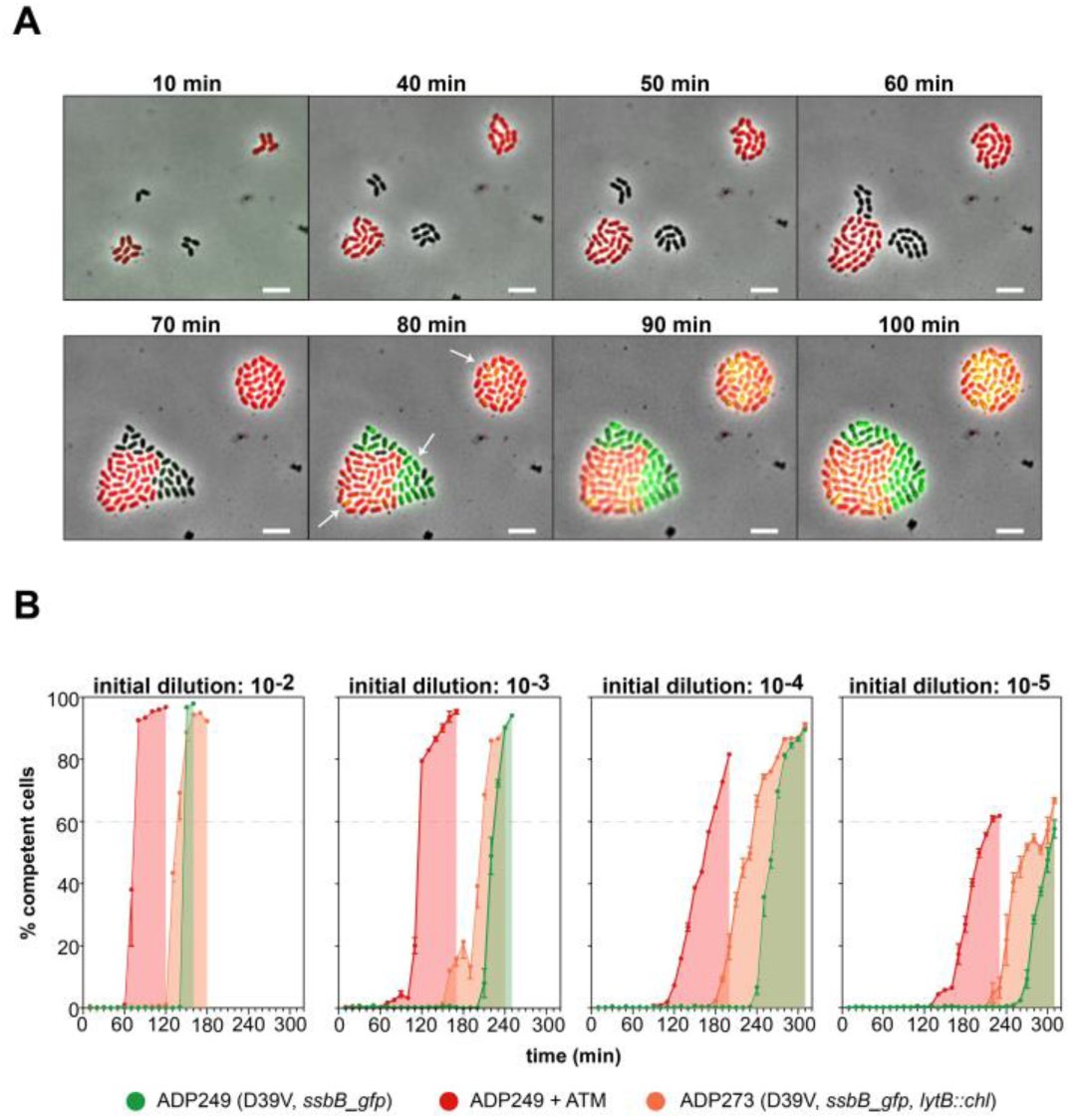
CSP is shared in a common pool and synchronizes initiation of competence. **A) Time-lapse fluorescence microscopy.** Two colonies with a fusion of the late competence gene *ssbB* to *gfp* are shown; one formed by cells of wild-type D39V (ADP249) and one formed by cells of a Δ*comC* mutant D39V (ADP247), which also constitutively expresses a red fluorescent protein. White arrows in the 80 minutes frame show that both D39V and Δ*comC* microcolonies became competent at the same time, independent of whether cells touch each other (left microcolony) or not (right microcolony). Scale bar: 4 μm. Note that the overlap, in the Δ*comC* strain, of green SsbB-GFP foci with the red background causes the foci to appear yellow. **B) Synchronization of competence at the single-cell level**. Cells of strains P_*ssbB*_-*ssbB*-*gfp* (ADP249) and P_*ssbB*_-*ssbB*-*gfp*, *lytB::chl* (ADP273) were grown in C+Y at competence-permissive pH 7.9; ADP249 was grown without antibiotics (green lines/areas) or with 28 μg/ml of aztreonam (red lines/areas), the Δ*lytB* ADP273 strain (orange lines/areas) without antibiotics. The highly permissive pH 7.9 used in this experiment allowed earlier competence development compared with the pH used in Figure 4, thereby reducing the required number of flow cytometry reads. The average of 3 replicates and Standard Error of the Mean (SEM) are plotted for each of four inoculation densities: OD_595nm_ 10^-2^, 10^-3^, 10^-4^ and 10^-5^. Twelve thousand individual particles (single cells, diplococci and/or chains) were detected for each replicate every 10 minutes along the experiment.

Next, we tested whether the results observed at the population level were reproducible in single-cell-level experiments. First, we established the noise level of false positive particles in flow cytometry using the Δ*comC* strain (which cannot become naturally competent), which turned out to be less than 1% of the cells (Figure S15). Interestingly, we observed a strong correlation between the detection of the first subpopulation of positive single cells via flow cytometry (2.5% and 4.1% of 12,000 cells per histogram in ATM and control conditions, respectively) and the first value of ≥ 100 RLUs in the plate reader, which was considered a positive signal for competence activation (Figure S16A). Similar results were obtained by fluorescence microscopy, ruling out the presence of an early, pre-existing subpopulation of competent cells below the detection limit of our flow cytometer or plate reader (Figure S16B).

For a better understanding of how competence is initiated and spread at the single-cell level in the wild type population, we studied untreated wild-type cells, ATM-treated wild-type cells and the Δ*lytB* mutant at four different inoculum sizes, analyzing 36,000 single particles every 10 minutes by flow cytometry (3 replicates of 12,000 particles), again using the SsbB-GFP reporter (Figure 5B). The single-cell flow cytometry data shows that competence development is density-dependent in all three conditions, rather than time-dependent. For instance, in the wild type, the onset of competence in cultures with an inoculum size of 10^-5^ was delayed by more than 2 hours relative to inoculums of 10^-2^ (green areas, Figure 5B). Note that the SsbB-GFP fusion is much more stable than the luciferase reporter used in plate reader assays, and that GFP-based assays, therefore, do not reflect the narrow window of transcriptional activity that is (more) visible in the corresponding luciferase assays.

As observed in the plate reader experiments, the presence of ATM (Figure 5B, red) and the deletion of *lytB* (Figure 5B, orange) both led to earlier competence development from all inoculation densities, compared to the control condition (Figure 5B, green); however, the synchronization of competent cells in the presence of ATM or absence of *lytB* is reduced compared to the control conditions. This is especially clear for the smaller inoculums, where cells had more time to form chains. Interestingly, the loss of synchronization in the presence of ATM is largely reversed by the exogenous addition of 100 nM CSP_1_ at the moment the first competent cells were detected (Figure S17), confirming that there is a large portion of live cells that did not sense enough CSP to develop competence in the absence of exogenous CSP_1_. Indeed, in the presence of ATM or a *lytB* deletion, synchronization of ≥ 60% of the population takes nearly twice as long as in control conditions, as measured from the first positive time-point (Figure S18). The addition of exogenous CSP_1_ in the presence of ATM nearly compensates for this loss in synchronization.

To test whether addition of CSP_1_ eliminates the differences observed in Figure 5B between wild-type and Δ*lytB* strain, we added three different concentrations of CSP_1_ (1nM, 10nM and 100nM) 60 minutes into the experiment. This time point is well before the onset of natural competence, so the ComD receptor is not produced at high levels or saturated yet. Indeed, for all three CSP_1_ concentrations, competence profiles of wild type and Δ*lytB* cells are nearly identical (Figure S19).

Altogether, these results show that the initiation of competence is density-dependent with CSP acting as a quorum sensing agent. However, this sensing can be disrupted or complicated by several factors, such as the presence of long chains retaining CSP, acidification of the medium by fermentation, or other phenomena that affect the diffusion of CSP into the common pool. Furthermore, once competence has initiated at lower cell densities, contact-dependent triggering of competence may play a role (Prudhomme et al., 2016) as exhibited by reduced propagation kinetics (Figure 5B).

### Cell chaining reduces the shared CSP pool

To elucidate whether production and export of CSP are affected in chaining cells, we employed the HiBiT tag detection system (Aggarwal et al., 2018; Wang et al., 2018). The HiBiT tag was placed under the control of the comCDE promoter, either with (strain ADP308) or without (strain ADP312) the leader peptide sequence of comC. As an additional control, we deleted comAB from strain ADP308 (strain ADP311). If the HiBiT peptide carries the leader sequence, it is recognized, cleaved and secreted by ComAB. Then, extracellularly, it reacts with a HiBiT-dependent luciferase variant (LgBiT), added in the medium, resulting in bioluminescence (Figure 6A, left). In the absence of the *comC* leader sequence, HiBiT accumulates in the cytoplasm and no luminescence is generated (Figure 6A, right). The extracellular bioluminescence produced by this reporter was similar in the wild type (ADP308) and the Δ*lytB* mutant (ADP310) (Figure 6B). We used strains ADP311 (Δ*comAB*) and ADP312 (no *comC* leader) to confirm that luminescence resulted from active export of the HiBiT tag and was not caused by cell lysis. In both strains, HiBiT cannot be exported and therefore accumulates in the cytoplasm. Indeed, although we detected some lysis after 120 minutes, the bioluminescence observed is significantly less compared to the strains that export the peptide (Figure 6B). Combined, these results strongly suggest that *comC* transcription and ComAB activity is not affected by cell chaining and CSP is exported at similar rates in chains of cells.

**Figure 6.**
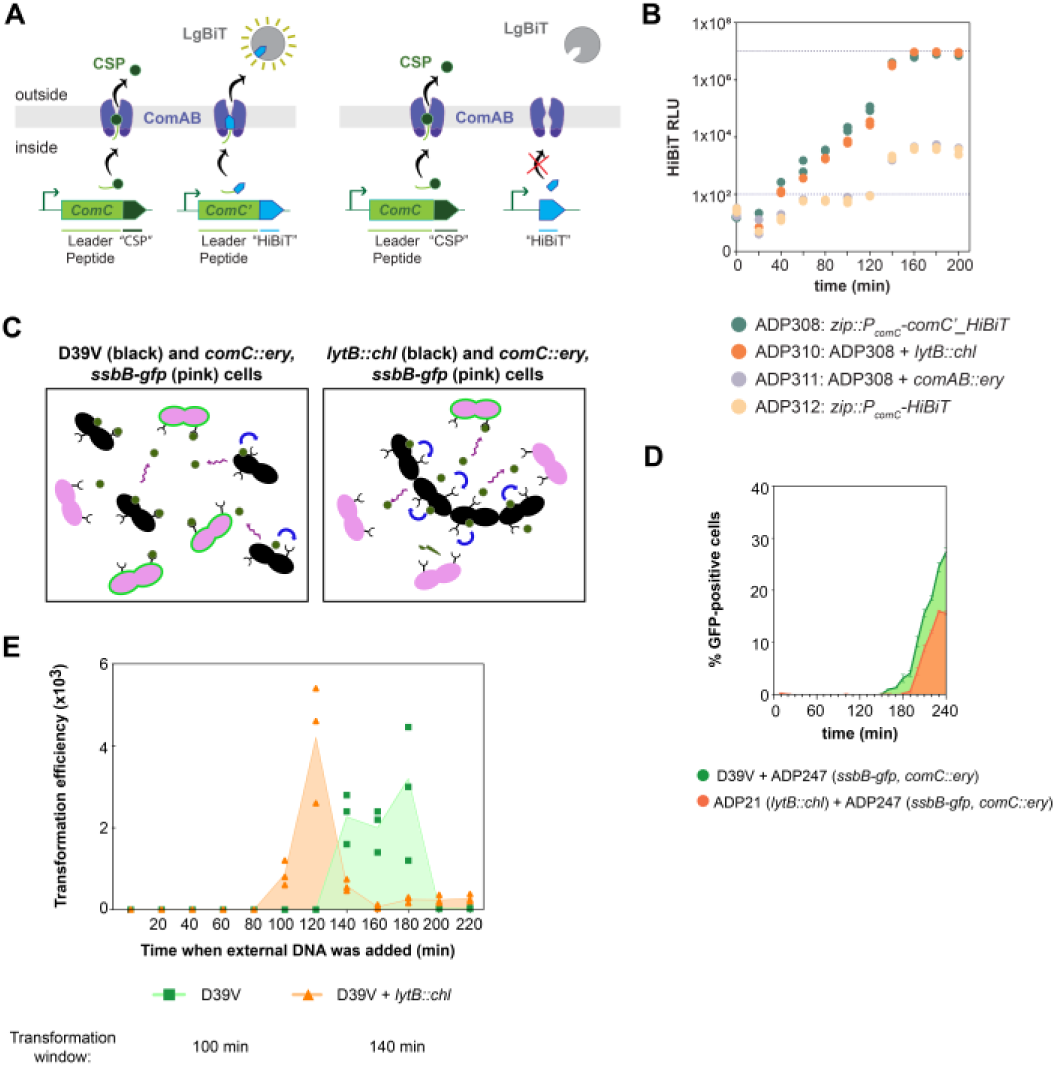
Cells in chains have similar CSP production levels but retain more CSP leading to an extended transformation period. **A) Graphical representation of the HiBiT experiment.** Left, ComC (called CSP once outside the cell) and HiBiT are regulated by the *comCDE* promoter, and both precursors have a leader peptide signal which is recognized, cleaved and exported by ComAB. Once outside the cell, HiBiT interacts with the soluble protein LgBiT and yields bioluminescence (Wang et al., 2018). Right, HiBiT lacks the ComC leader peptide and accumulates in the cytoplasm since it cannot be recognized and exported by ComAB. **B) CSP is exported at a similar rate in wild type and Δ*lytB* mutant cells.** Bioluminescence (Relative Luminescence Units, RLU) can be correlated with CSP export. In both the wild type (ADP308) and the Δ*lytB* mutant (ADP310), the export rates are similar until the saturation point (1×10^7^ RLU). Cells were grown in C+Y at competence-permissive pH 7.6. At pH 7.6, cells become naturally competent but with a delay relative to pH 7.9, facilitating the visualization of the inducing effect of chaining. However, competence development occurs later than the RLU saturation point. Neither the *comAB* mutant (ADP311) nor the HiBiT version without the leader peptide (ADP312) showed any signal during the first 120 min (values below the threshold line of 100 RLU). After that, potentially due to cell lysis, the signal increased but was negligible compared to the exported version of the peptide. Two replicates are shown for each time point and condition. **C) Graphical representation of the experimental setup.** Coincubation (1:1 proportion) of wild type D39V (black, left panel) or *ΔlytB* (black, right panel) with *ΔcomC* mutant (pink) cells. D39V releases more CSP (green dots) into the common pool than the Δ*lytB*, and more Δ*comC* cells become competent (green halo). **D) Competence induction in a *comC* mutant by wild-type D39V and the *lytB* mutant.** Strains were coincubated (1:1) in C+Y medium (pH 7.9), with an initial density OD_595nm_ of 10^-4^. Three replicates were analyzed by flow cytometry every 10 minutes to detect GFP signal (12,000 particles each). Green: coincubation of D39V and ADP247 (D39V, *ssbB*-*gfp*, *comC::ery*); orange: coincubation of ADP21 (*lytB::ch1*) and ADP247. Note that on the Y-axis the percentage of GFP-positive cells reflects the percentage of all cells in the population (including cells not harboring the SsbB-GFP fusion). **E) Cell chaining widens the transformation time window.** D39V (green) and *ΔlytB* (orange) were grown in C+Y at pH 7.9. Every 20 minutes, 1 μg of naked linear DNA with homology regions of 1 kb around the *ssbB* locus (*ssbB*-*luc*-*kan*, see M&M) was added. After 20 minutes, the cultures incubated with DNA were plated with and without 250 μg/ml kanamycin, to collect the number of transformants and total viable counts, respectively. Three replicates are plotted for each condition. The highlighted area shows the temporal window in which cells were able to take up DNA.

As the amount of CSP released is similar in wild-type and Δ*lytB* cultures, we believe that the chain-induced phenotype retains CSP and decreases the amount of CSP released to the shared pool, reducing the synchronization of the population (Figure 6C). Thus, chain formation would reshape global quorum sensing signaling, where all cells communicate and synchronize competence in a short lapse of time, into local quorum sensing signaling, where chains retain and sense most of their own produced CSP. To test this hypothesis, we analyzed the ability of wild-type D39V and the Δ*lytB* strain to induce competence in a coincubated Δ*comC* strain that harbors the SsbB-GFP fusion. The Δ*comC* strain is only able to become competent if there is free CSP in the medium, but cannot produce its own CSP. As shown in Figure 6D, competence in the Δ*comC* strain was detected roughly 40 minutes earlier when mixed with wild type cells than with the Δ*lytB* mutant. This seems in contrast with the fact that the *ΔlytB* strain is hypercompetent and therefore should release CSP into the medium earlier than the wild type (in individual populations, the Δ*lytB* mutant became competent 60 minutes earlier than the wild type; Figure 5B). Furthermore, the fraction of activated Δ*comC* cells incubated with wild-type cells was nearly twice as high as for cells coincubated with Δ*lytB* (Figure 6D). The initial presence of chains was prevented by bead-beating and there was no significant difference in either growth rate or survival rate between the Δ*lytB* and wild type. Therefore, these results support the conclusion that wild-type D39V releases more CSP into the common pool than the Δ*lytB* mutant, leading to earlier competence activation in the Δ*comC* strain.

Finally, we studied the effect of CSP concentration on the synchronicity of competence development throughout the population. To this end, we added different concentrations of exogenous synthetic CSP_1_, either at the beginning of the experiment or 90 minutes after, just before the onset of competence (Figure S20). Interestingly, the dynamics of competence propagation are similar for different CSP concentrations, with a concentration-dependent delay in the onset of competence. When CSP_1_ was added after 90 minutes (roughly three doubling times), this delay in offset was not visible. The dynamics of propagation were similar, with more than 60% of the population becoming competent 20 minutes after the addition of CSP_1_. Together, this data shows that chained pneumococci have distinctly different kinetics of competence activation and signal propagation from unchained, untreated wild-type diplococci, and do not contribute as much to the extracellular pool of CSP.

### Natural competence in chained bacteria extends the transformation window

To investigate the biological relevance of the chain-induced phenotype, we performed transformation experiments, adding external DNA every 20 minutes in the D39V and Δ*lytB* strains (Figure 6E). The presence of a chaining phenotype increases the window where bacteria can take up and integrate exogenous DNA, from 100 min (in D39V) to 140 min (in Δ*lytB*) in our set-up.

### Reduced diffusion stimulates competence development

Since competence is initiated by recognition of extracellular CSP by its cognate membrane-anchored receptor (ComD), on the basis of the data presented so far, we speculated that chain formation alters CSP diffusion and thereby sensing by ComD. To test whether a reduction of CSP diffusion in the medium can also trigger and desynchronize competence, we decreased the diffusion coefficient of the growth medium by making it more viscous. To do so, we grew cells in a range of concentrations of Pluronic-127, an innocuous polymer that increases the medium density without affecting cell metabolism (Yang et al., 2010). Indeed, pneumococcal cell morphology and cell chain length are unaltered by the addition of Pluronic-127 during exponential growth (Figures S10 and S21). In line with our model, the addition of this polymer resulted in earlier competence development (Figure S21A). However, we observed a reduced slope of *P_ssbB_*-controlled luciferase expression (RLU) at several inoculation densities, at the Pluronic-127 concentrations tested (Figure S21B).

Although the exponential growth rates in the presence or absence of multiple concentrations of Pluronic-127 were identical, indicating a normal cell metabolism, we performed time-lapse microscopy to confirm that this polymer does not promote cell clumping or biofilm-like phenotypes. Using the same set up as for Figure 5A, we observed that in spite of the increased viscosity in the medium, some CSP can still diffuse in the medium and activate the Δ*comC* cells. This activation occurs without direct cell-to-cell contact (Figure S22). The normal phenotype of cells in presence of Pluronic-127 (Figure S10), together with the loss of synchronization (Figure S21B), suggests that reducing CSP diffusivity is the dominant effect of this polymer. Even if we cannot discard that Pluronic-127 has another effect inducing competence, these results confirm that its main role is decreasing the diffusion of CSP_1_, which results in an upregulation of competence. Together, these data suggest that reduced diffusion of the CSP pheromone in the medium results in earlier activation of local CSP-ComD-ComE positive feedback loops, which at the same time desynchronizes population-wide competence activation.

## Discussion

Many clinically used antibiotics are able to induce competence, which can subsequently lead to the acquisition of antibiotic resistance. Two molecular mechanisms underlying antibiotic-induced competence have been described: altered gene dosage by DNA-targeting antibiotics (Slager et al., 2014), and reduced CSP degradation by HtrA under mistranslation conditions (Stevens et al., 2011). The principal contribution of this work is the identification of a third mechanism, by which certain cell-wall-targeting antibiotics can induce competence. Specifically, the antibiotic aztreonam (ATM), which is used to treat respiratory infections caused by Gram-negative bacteria, and clavulanic acid (CLA), which is frequently co-administered with the broad-spectrum antibiotic amoxicillin, induce competence (Figure 1B).

Both ATM and CLA target the non-essential PBP3 of *S. pneumoniae* (Figure 3B) (Kocaoglu et al., 2015; Severin et al., 1997), and we show that this causes cell chaining (Figures S9 and 3D). Using CRISPRi-mediated depletion of *pbp3* and deletion of the major autolysin LytB, known to result in long chains, we confirmed that competence is upregulated when pneumococci form chains instead of having the normal diplococcal appearance (Figures 3A and S13).

It is interesting to note that our observations reconcile observations made across different laboratories concerning the dynamics of pneumococcal competence. For instance, at a single-cell level we confirmed that competence occurs first in a small subpopulation, and then spreads to the whole population, as suggested by Prudhomme and coauthors (Prudhomme et al., 2016). However, the way in which competence is propagated still remains a cause of debate; some evidence indicates that CSP is, to some extent, retained by producer cells and competence propagates by cell-cell contact (Prudhomme et al., 2016; Fig. 7). However, other data showed that CSP is released into a common, shared pool and sensed by the whole population in a typical quorum sensing manner, which does not require direct contact between cells (Fig. 5A; Moreno-Gámez et al., 2017). Here, we show that despite the appearance of a small initial subpopulation of competent cells, in normal conditions (diplococcal phenotype), competence is rapidly spread and synchronized (Figures 4 and 5B). Under cell chaining conditions or when the medium acidifies after several hours of cultivation, the dynamics of competence propagation seems to depend more on short-range communication between cells (Figure 5B). However, the similar dynamics of population-wide competence development in the presence of various concentrations of exogenous CSP_1_, even in the presence of ATM, supports the existence and importance of a quorum sensing mechanism, in addition to a contact-dependent mechanism of competence propagation (Figures S17 and S19).

**Figure 7.**
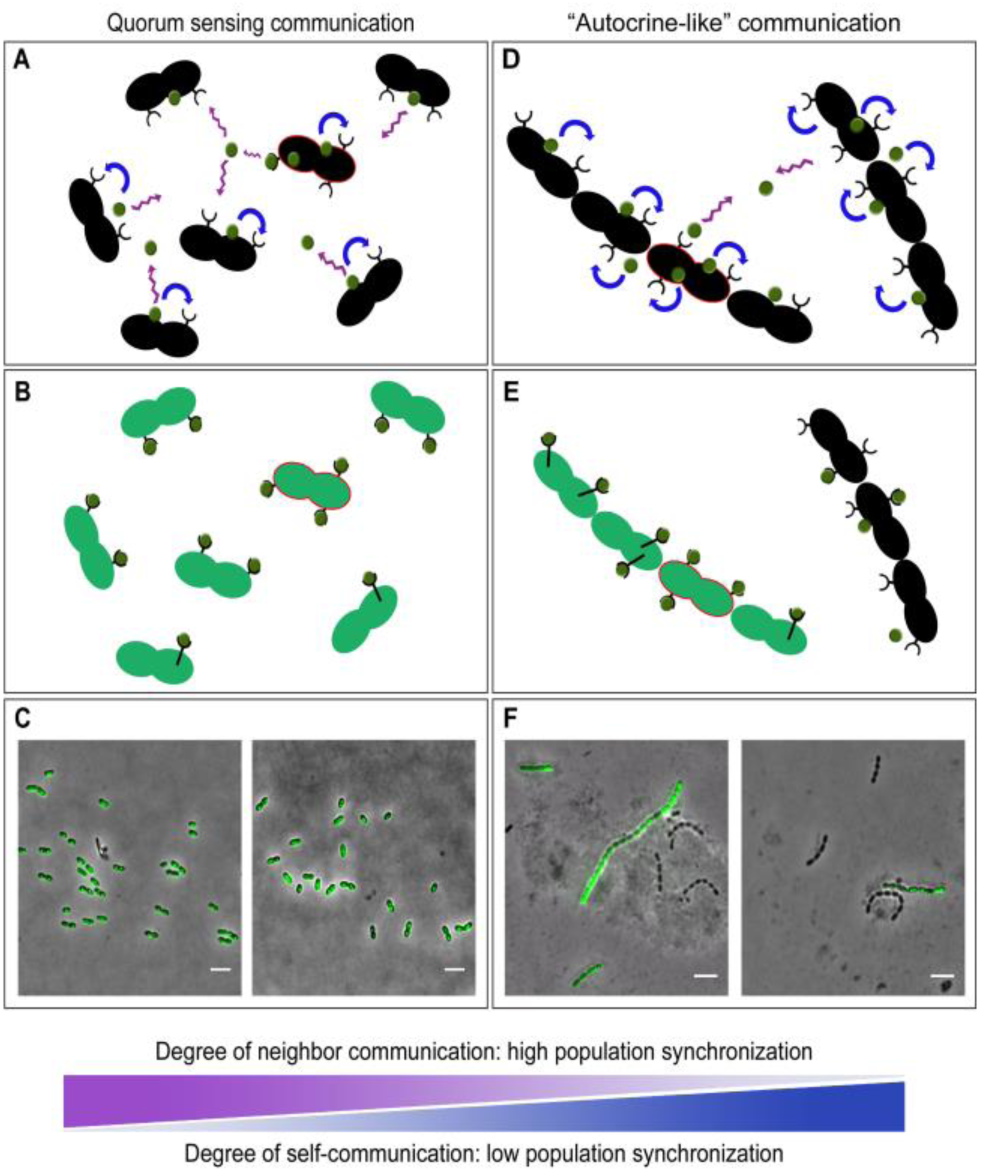
Models of global quorum sensing signaling (left) and local quorum sensing or autocrine-like signaling (right). **A)** *S. pneumoniae* secretes CSP to communicate with other cells (purple arrows) and to synchronize competence once a critical CSP (green dots) threshold is reached. In addition, self-sensing CSP (blue arrows) plays a role, as part of the CSP pool is retained by the producer cell (Prudhomme et al., 2016). Diplococci producing more CSP than average (highlighted in red) contribute to the increase of the extracellular CSP pool. **B)** Because of neighbor communication, when the CSP threshold is reached, all diplococci synchronize and become competent at nearly the same time (green cells). **C)** Microscopy of the ADP249 strain (*P_ssbB_*-*ssbB*-*gfp*) after the first competence time point, showing that all diplococci synchronized and expressed GFP at the same time. When the first GFP signal was observed in the plate reader, cells were collected and 100 μg/ml of chloramphenicol (to stop translation) and 4% paraformaldehyde (to fix the cells) were added. Cells, then, were resuspended in 5 μl of PBS to concentrate them and imaged by fluorescence microscopy. Scale bar: 4 μm. Two different fields of view are shown. **D)** Chain formation shifts quorum sensing signaling from a global mechanism to a local mechanism. CSP released by cells present in the same chain is retained and sensed by the same chain. **E)** As a result of this local quorum sensing, the extracellular pool is smaller, thereby reducing the communication with other cells and decreasing competence synchronization (not all the cells become competent at the same time). However, as stochastic fluctuations are not buffered through the shared pool of CSP, individual chains of cells will initiate competence earlier than well-mixed populations consisting of diplococci. **F)** Microscopy of the ADP273 strain (*lytB::chl*, *P_ssbB_*-*ssbB*-*gfp*) after the first competence timepoint, showing that not all the chains expressed GFP as a result of the disruption of competence signaling. Images were taken as in panel C.

Interestingly, coincubation of the Δ*comC* reporter strain together with either wild-type D39V (diplococcal phenotype) or Δ*lytB* (chained phenotype) cells revealed that diplococci more efficiently induce competence in the Δ*comC* mutant (Figure 6D). This evidence is even more marked if we consider that the Δ*lytB* is hypercompetent (Figure 5B) and produces CSP earlier in time, relative to the D39V cells. These results strongly suggest that when pneumococci grow as diplococci, the produced CSP is mostly shared in the medium and sensed by >60% of the Δ*comC* cells (Figure 6D), which cannot become naturally competent but can sense the exogenous CSP released by the wild type. Notably, we demonstrated in a time-lapse microscopy experiment that the activation of competence in the Δ*comC* mutant does not require direct contact with the producer (Figure 5A). In contrast, the presence of long chains of producer cells results in a delayed competence induction of Δ*comC* cells, and only 30% of the Δ*comC* population sensed enough CSP to activate competence (Figure 6D). As the growth rate, the initial number of cells and the CSP production rate are the same in D39V and the Δ*lytB* strain (Figure 6, De las Rivas et al., 2002; García P et al., 1999), the differences between both conditions can only be explained by a retention of the CSP by the chain-induced phenotype, thereby limiting the diffusion of CSP into the common pool.

At the population and single-cell level, this creates local positive-feedback loops that result in a competence response that starts earlier, but is less synchronized (Figures 4 and 5). In addition, the presence of chains could decrease the local or global diffusivity of the CSP in the medium, enhancing this local quorum sensing signaling (Figure S21B).

Pneumococcal competence is a population sensing process that, to a certain extent, is influenced by stochastic parameters, such as basal ComAB and ComCDE expression, replication state and many more indirect factors. Therefore, single cells produce and sense CSP at different rates and differences in local CSP concentration will occur. These differences, along with heterogeneity in cells’ CSP-sensing potential will lead to slight timing differences of competence activation on a single-cell level, thereby leading to the formation of initial subpopulations of competent cells that then activate the rest of the population. Also, competent cells produce cell wall hydrolases and might reduce growth and kill non-competent siblings (Claverys et al., 2007). Interestingly, several factors, such as pH or antibiotics, can modify the rates at which single cells produce and/or sense CSP (Moreno-Gámez et al., 2017; Prudhomme et al., 2016). Our results suggest that chain formation by the presence of ATM or CLA modifies the balance between CSP production and sensing, increasing the self-sensing of CSP between cells within the same chain. Thus, single cells that produce more CSP than average are more likely to share this CSP with cells of the same chain (autocrine-like signaling), reducing the shared pool of CSP (Figure 7, right) (Bareia et al., 2017).

We propose to keep using the term quorum sensing (QS) to describe competence activation and signal propagation, as it is clear in the field, as nicely stated by Paul Williams “that the size of the ‘quorum’ is not fixed but depends on the relative rates of production and loss of the signal molecule, which will, in turn, vary depending on the local environmental conditions” (Williams P, 2007). In addition, Williams also pointed out that QS can also be considered in the context of “diffusion or compartment sensing”, where the signal molecule supplies information with respect to the local environment and spatial distribution of the cells rather than, or as well as, “global cell population density” (Williams P, 2007). This beautifully sums up the observations made here for competence development in *S. pneumoniae*.

Notably, other clinical beta-lactam antibiotics such as aminopenicillins or cephalosporins are not able to induce competence as they do not bind PBP3, but PBP2B and/or PBP2X (Kocaoglu et al., 2015). Contrary to PBP3, the depletion of PBP2B and PBP2X does not lead to chain formation, and thereby to upregulation of competence (Figures 3A and S10).

Amoxicillin/clavulanic acid (Augmentin) has been available for over 20 years and continues to be one of the most widely used antibiotics, especially in the treatment of respiratory tract infections. However, CLA is a beta-lactamase inhibitor that is useless for the specific treatment of pneumococcal infections, as there have been no reports of *S. pneumoniae* producing beta-lactamases. Our study suggests that in such cases clavulanic acid can best be omitted for antibiotic therapy as it would drive pneumococcal evolution and potentiate antibiotic resistance development by upregulating competence.

Additionally, it has been described that the presence of pneumococcal chains enhances adhesion and colonization (Rodriguez et al., 2012), facilitating the persistence in the nasopharynx in pneumococcal (or polymicrobial) biofilms. This chained phenotype could result in a prolonged time window, during which cells are able to take up exogenous DNA (Fig. 6E), and explain the rapid adaptation and evolution in response to antibiotic-induced stress in pneumococcal strains colonizing the nasopharynx (Croucher et al., 2011). Thus, it will be interesting to see how competence is synchronized and propagated in more realistic environments, closely resembling the polymicrobial environment that is present in the human nasopharynx. Continued molecular epidemiology studies will be crucial to determine the role and long-term effects of antibiotic therapy and vaccination on pneumococcal prevalence and antibiotic resistance.

## Materials and Methods

### Bacterial strains and growth conditions

All pneumococcal strains used in this study are derivatives of the clinical isolate *S. pneumoniae* D39V (Avery et al., 1944; Slager et al., 2018) unless specified otherwise. See Table S6 for a list of the strains used and the Supplemental information for details on the construction of the strains.

*S. pneumoniae* was grown in C+Y medium at 37°C. C+Y was adapted from Adams and Roe (Adams & Roe 1945) and contained the following compounds: adenosine (68.2 μM), uridine (74.6 μM), L-asparagine (302 μM), L-cysteine (84.6 μM), L-glutamine (137 μM), L-tryptophan (26.8 μM), casein hydrolysate (4.56 g L^-1^), BSA (729 mg L^-1^), biotin (2.24 μM), nicotinic acid (4.44 μM), pyridoxine (3.10 μM), calcium pantothenate (4.59 μM), thiamin (1.73 μM), riboflavin (0.678 μM), choline (43.7 μM), CaCl_2_ (103 μM), K_2_HPO_4_ (44.5 mM), MgCl_2_ (2.24 mM), FeSO_4_ (1.64 μM), CuSO_4_ (1.82 μM), ZnSO_4_ (1.58 μM), MnCl_2_ (1.29 μM), glucose (10.1 mM), sodium pyruvate (2.48 mM), saccharose (861 μM), sodium acetate (22.2 mM) and yeast extract (2.28 g L^-1^).

We can control competence development by changing the pH in the medium. The underlying mechanism it is not fully understood, but it is believed that is related to the production and export of CSP (Moreno-Gámez et al., 2017). For this reason, we always grow a preculture in C+Y at pH 6.8, because at this pH, even the hypercompetent strains such as Δ*lytB* or Δ*pbp3* mutants, are not able to accumulate enough CSP to induce competence before cells reach stationary phase.

### Luminescence assays of competence induction

To monitor competence development, strains either contain a transcriptional fusion of the firefly *luc* and the *gfp* gene with the late competence gene *ssbB* or a full translational *ssbB*-*gfp* fusion. Cells were pre-cultured in C+Y (pH 6.8) at 37°C to an OD_595nm_ of 0.4. Right before inoculation, cells were collected by centrifugation (8000 rpm for 3 minutes) and resuspended in fresh C+Y at pH 7.3, which is non-permissive for natural spontaneous competence under these experimental conditions. All experiments were started with an inoculation density of OD_595nm_ 0.004, unless indicated. Luciferase assays were performed in 96-wells plates with a Tecan Infinite 200 PRO illuminometer at 37°C as described before (Slager et al., 2014). Luciferin was added at a concentration of 0.45 mg/mL to monitor competence by means of luciferase activity. Optical density (OD_595nm_) and luminescence (relative luminescence units [RLU]) were measured every 10 minutes. For the CRISPRi experiments, cells were grown as above, and diluted 100x in the presence of a range of IPTG indicated for each condition, depending on whether the targeted gene is essential or not. Despite the fine-tuning regulation of CRISPRi, there is some leakiness that could slightly affect the growth rates and time of natural competence development. For this reason, in these experiments, we do not compare the effect between strains but we compare the control with the addition of IPTG in every strain.

### Detection of the PBPs using Bocillin-FL

Samples were prepared as described before (Kocaoglu et al., 2015) with slight modifications. Briefly, 4 ml of cells were grown in C+Y pH 6.8 until OD 0.15 and harvested by centrifugation (16,000 × *g* for 2 min at 4 °C). Cell pellets were washed in 1 ml PBS, pH 7.4. Cells were pelleted and resuspended in 50 μl PBS with or without the indicated concentration of ATM or CLA. After 30 min of incubation at room temperature, cells were pelleted, washed in 1 ml PBS, and resuspended in 50 μl PBS containing 5 μg/ml Bocillin-FL. After 10 min of incubation at room temperature, cells were washed again in 1 ml PBS. Next, cells were sonicated on ice (power 30%, three cycles of 10 seconds interval with a 10 seconds cooling time on ice (Sonoplus, Bandelin). Then samples were centrifuged at max speed for 15 min at 4°C and pellets were resuspended in 100 μl cold PBS. The protein concentration was adjusted to 2 mg/ml as determined by Bradford by diluting with PBS. 5x SDS-PAGE loading buffer was added to each sample and heated 10 minutes at 95 °C. Proteins were separated by gel electrophoresis (10% acrylamide) for 2.5 h at 180 V, 400 mA, and 60 W. The gel was scanned using a Typhoon gel scanner (Amersham Biosciences, Pittsburgh, PA) with a 526-nm short-pass filter at a 25-μm resolution.

### Intraspecies horizontal gene transfer (HGT)

We calculated the *in vitro* HGT efficiency using two genetically identical pneumococcal strains, differing only with the integration of two antibiotic resistance markers at two different locations of the genome. Strains DLA3 and MK134 (tetracycline and kanamycin resistant, respectively), (Slager et al., 2014) were grown to OD_595nm_ 0.4 in C+Y pH 6.8 at 37°C (non-permissive conditions for natural competence activation). Then, a mixed 100-fold dilution of both strains were grown in C+Y pH 7.3 (non-permissive conditions) and pH 7.5 (permissive conditions) to OD_595nm_ to promote the transfer of genes. When cells reached OD_595nm_ 0.4 again (approximately 3 hours), serial dilution of cultures were plated in Columbia agar + 5% sheep blood with 250 μg/ml of kanamycin plus 1 μg/ml tetracycline for the recovery of the number of recombinants, and without antibiotics to obtain the total viable counts, respectively) . Plates were incubated for 16h at 37°C with 5% CO2.

### Interspecies horizontal gene transfer

*S. pneumoniae* strain D39V was grown to OD_595nm_ 0.4 in C+Y pH 6.8 at 37°C, and *E. coli* carrying the plasmid pLA18 (integrates the tetracycline resistant marker *tetM*, via double crossover at the non-essential *bgaA* gene in *S. pneumoniae*, and contains a high copy Gram-negative origin of replication; Slager et al., 2014) was grown overnight with shaking, in LB supplemented with 100 μg/ml of ampicillin (resistant marker also contained in the plasmid, outside the double integration region). Both strains were diluted to OD_595nm_ 0.004 and co-incubated with or without 28 μg/ml of ATM in C+Y pH 7.3. After 3h, serial dilutions were plated either with 1 μg/ml of tetracycline (to recover transformants) or 50 μg/ml of aztreonam (to recover only the total viable pneumococci). Transformation efficiency was calculated by dividing the number of transformants by the total number of viable count. Three independent replicates of each condition were performed.

### Microarray experiments

Pneumococcal transcriptome profiles in the presence or absence of antibiotics were tested under conditions that do not support natural competence development to avoid differences in gene expression due to the activation of the competence pathway. We used strain *S. pneumoniae* ADP62 (D39V non-competent variant, *comC*::*ery*), grown in two biological replicates in C+Y (pH 7.6). Two kind of experiments were performed to detect rapid and adaptive exposures to the antibiotics. For the fast response, cells were collected during mid-exponential growth phase (OD 0.15) and incubated 15 minutes with or without 2 μg/ml of CLA or 28 μg/ml of ATM. For the adaptive response, cells at OD 0.15 were diluted 100X with or without the same concentration of antibiotics and grown again until OD 0.15. Results were compared using DNA microarray analysis, as previously described. (Shafeeq et al., 2015). For the identification of differentially expressed genes a Bayesian p< 0.001 and a fold change cut-off ≥2 was applied. Microarray data are available at Gene Expression Omnibus (GEO) with accession number GSE111562.

### oriC-ter ratio determination by Real-Time qPCR

Cells were grown as described above in the presence of antibiotics. In the real-time qPCR experiments, samples were prepared as previously detailed (Slager et al., 2014). Amplification was performed on a iQ5 Real-Time PCR Detection System (Bio-Rad). Amplification efficiencies and analysis were performed as before (Slager et al., 2014).

### Chain formation detection

To detect morphological changes, we incubated the different strains in C+Y acid medium (pH 6.8) until OD_595nm_ 0.1 and OD_595nm_ 0.4. Antibiotics or IPTG were added when indicated. 1 μl of cells at the indicated optical density was spotted onto a PBS agarose pad on microscope slides, and phase contrast images were acquired with a Leica DMi8 microscope. Microscopy images conversions were done using Fiji and analysis of the length of the chains was done using MicrobeJ (Ducret et al., 2016). Plotting was performed using the BactMAP/spotprocessR package (Van Raaphorst R, in preparation; https://github.com/veeninglab/spotprocessR).

### Fluorescence microscopy

To detect the morphological changes after incubation with antibiotics, 1 μl of cell suspension was spotted onto a PBS agarose pad on microscope slides. Phase contrast images were acquired with a Leica DMi8 microscope with a DFC9000 GT camera and a 100x/1.42 NA phase/c lens. Images were analyzed with ImageJ. For fluorescence microscopy of strains containing SsbB-GFP fusions, cells were spotted onto agarose slides as detailed above, and visualization was performed using a SpectraX light engine (Lumencor) using the following filters for GFP: Quad mirror (Chroma #89000), excitation at 470/24 nm, emission at 515/40 nm. For mKate2 (RFP): Chroma #69008 with excitation at 575/35 nm and emission at 600-670.

Time-lapses videos were recorded by taking images every 10 minutes. The polyacrylamide gel used as semi-solid growth surface was prepared with C+Y (pH 7.9) and 10% acrylamide.

### Flow cytometry

ADP245 (P_*ssbB*_-*ssbB*-*gfp*, *bgaA::*P_*ssbB*_-*luc*) or ADP249 cells (P_*ssbB*_-*ssbB*-*gfp*) cells were pre-cultured in C+Y (pH 6.8) at 37°C to an OD_595nm_ of 0.1, washed and diluted as explained before in C+Y (pH 7.9). Cells were thoroughly vortexed to avoid possible chains. Experiments were started with an inoculation density of OD_595nm_ 0.0001, with or without 28 μg/ml of ATM. Optical density (OD_595nm_) was measured every 10 minutes in 96-wells plates with a Tecan Infinite 200 PRO luminometer at 37°C. Right after every measurement, a sample was taken and measured on a Novocyte Flow Cytometer (ACEA Biosciences). The pneumococci were gated to exclude debris. Twelve thousand bacteria were analyzed for FITC fluorescence using a 488 nm laser (GFP expression) with a flow rate of 9 μl/min. Cells pretreated with CSP_1_ and cells untreated were used to establish the cutoff value for FITC positive (competence activation). Results were analyzed by Novoexpress software (ACEA Biosciences).

### Nano-Glo HiBiT Extracellular Detection System

Cells were pre-cultured in C+Y (pH 6.8) at 37°C to an OD_595nm_ of 0.1, washed and diluted as explained before in C+Y (pH 7.6). Experiments were started with an inoculation density of OD_595nm_ 0.001. Optical density (OD_595nm_) was measured every 10 minutes in 96-wells plates with a Tecan Infinite 200 PRO luminometer at 37°C. Every 20 minutes, 50 μl of the Nano-Glo Extracellular Detection System reagent was added as specified in the manufacturer’s instructions. Additionally, media and PBS samples were used as controls. Bioluminescence was measured every minute during the 10 minutes after reagent addition.

### Quantification and statistical analysis

Data analysis was performed using GraphPad Prism and Microsoft Excel. A one-tailed student t test was used to determine differences on chain formation and transformation efficiency.

Data shown in plots are represented as mean of at least three replicates ± SEM, as stated in the figure legends. Exact number of replicates for each experiment are enclosed in their respective figure legends.

## Footnotes

**Author Contributions and Notes** A.D. and J.W.V. designed research, A.D. and J.S performed research, A.D. and J.S. analyzed data; and A.D. and J.W.V. wrote the paper. The authors declare no conflict of interest. This article contains supporting information online.

## Acknowledgments

Work in the Veening lab is supported by the Swiss National Science Foundation (project grant 31003A_172861), a VIDI fellowship (864.11.012) of the Netherlands Organization for Scientific Research, a JPIAMR grant (50-52900-98-202) from the Netherlands Organisation for Health Research and Development (ZonMW) and ERC starting grant 337399-PneumoCell. A.D. was supported by Marie Skodowska-Curie fellowship 657546.

